# High prevalence of Prdm9-independent recombination hotspots in placental mammals

**DOI:** 10.1101/2023.11.17.567540

**Authors:** J. Joseph, D. Prentout, A. Laverré, T. Tricou, L. Duret

## Abstract

In many mammals, recombination events are concentrated into hotspots directed by a sequence specific DNA-binding protein named Prdm9. This protein facilitates chromosome pairing and its inactivation has been shown to induce fertility losses in mice and rats. Intriguingly, *Prdm9* has been lost several times in vertebrates, and notably among mammals, it has been pseudogenized in the ancestor of canids (dogs, wolves foxes). When this gene is inactive, either naturally in dogs, or through knock-out experiments in mice, recombination hotspots still exist, but they tend to occur in promoter-like features such as CpG islands. It has thus been proposed that one role of *Prdm9* could be to direct recombination away from those Prdm9-independent hotspots. However, the ability of Prdm9 to direct recombination hotspots has been assessed only in a handful of species, and a clear picture of how much recombination occurs outside of Prdm9-directed hotspots in mammals is still lacking. In this study, we derived an estimator of past recombination activity based on signatures of GC-biased gene conversion in substitution patterns. We applied it to quantify recombination activity in Prdm9-independent hotspots in 52 species of boreoeutherian mammals. We observed a wide range of recombination rate at these loci: several species (such as mice, humans, some felids or cetaceans) show a deficit of recombination, while a majority of mammals display a clear peak of recombination. Our results demonstrate that Prdm9-directed and Prdm9-independent hotspots can co-exist in mammals, and that their co-existence seem to be the rule rather than an exception.

## Introduction

Meiotic recombination is a crucial step in the production of gametes for a large majority of eukaryotes. It is initiated by programmed double-strand breaks (DSBs), that can be resolved in two types of recombination events, using the homolog as a template: crossovers (COs), where there is a reciprocal exchange of chromosome arms, and non crossovers (NCOs), where DSB repair leads to gene conversion around the DSB site without reciprocal exchange. In most eukaryotes, at least one CO per chromosome is necessary to ensure correct segregation between homologs and is therefore mandatory for meiosis success (Page and Hawley, 2003; Gerton and Hawley, 2005). In many vertebrates, recombination events are not uniformly distributed along the genome (reviewed in Stapley et al. (2017); Zelkowski et al. (2019)). Instead, they tend to be concentrated in so-called recombination hotspots (Lichten and Goldman, 1995; Tock and Henderson, 2018).

In many mammals, the position of recombination hotspots is determined by the zinc-finger protein Prdm9, which binds specific DNA motifs and recruits the DSB machinery through histone methylation (Baudat et al., 2010; Myers et al., 2010; Parvanov et al., 2010; Diagouraga et al., 2018). This gene is highly polymorphic and hundreds of alleles have been reported in mice and humans (Buard et al., 2014; Kono et al., 2014; Alleva et al., 2021). Most allelic diversity is concentrated on residues of the zinc fingers that interact with the DNA, leading to changes in DNA sequence specificity. Therefore, the position of recombination hotspots varies within a population and between species (Auton et al., 2012; Smagulova et al., 2016; Alleva et al., 2021). Additionally, Prdm9 tends to erode its targets through gene conversion (Baker et al., 2015; Smagulova et al., 2016). As the available targets for a given Prdm9 allele become scarce, its ability to generate enough COs for meiosis to succeed is compromised. A new allele with more targets will then be positively selected leading to a red-queen dynamic accelerating the turnover of recombination hotspots (Úbeda and Wilkins, 2011; Latrille et al., 2017; Baker et al., 2022; Genestier et al., 2023). It has also been proposed that when Prdm9 binds symmetrically both homologs, it facilitates the repair of DSBs as COs and thereby contributes to the success of meiosis (Davies et al., 2016; Li et al., 2019; Hinch et al., 2019). Indeed, experiments in mice and rats showed that an inactivation of this protein drastically reduces fertility (Mihola et al., 2019, 2021; Brick et al., 2012).

Despite its central role in recombination, *Prdm9* has been repeatedly lost in vertebrates. (Baker et al., 2017; Cavassim et al., 2022). Among amniotes, one loss occurred in the ancestor of archosaurs (crocodiles and birds) (Singhal et al., 2015; Cavassim et al., 2022), and another one in the ancestor of canids (dogs, wolves and foxes) (Oliver et al., 2009; Axelsson et al., 2012; Auton et al., 2013). In dogs and several passerines, studies based on linkage disequilibrium (LD) showed that recombination hotspots tend to occur in CpG islands, and more specifically in hypomethylated ones in dogs (Auton et al., 2013; Berglund et al., 2015; Singhal et al., 2015; Kawakami et al., 2017). Likewise, the DSB hotspots of a mouse whose *Prdm9* has been inactivated through a knock-out (PRDM9^-/-^) also occur in promoter-like features (Brick et al., 2012). Interestingly, an increase of recombination rate near promoters has also been observed in plants and yeasts, which lack *Prdm9* (Petes, 2001; Marand et al., 2017).

Those observations led to the conclusion that there exist two types of recombination landscapes in verte-brates. The first one is Prdm9-dependent, fast evolving with recombination targeted away from functional elements, while the second is relatively stable, with recombination occurring in promoter-like feature such as CpG islands. However, a recent finding in snakes challenged this binary view. In rattle snakes and corn snakes, which still possess a functional *Prdm9*, some hotspots are directed by Prdm9, but others are directed toward CpG islands (Schield et al., 2020; Hoge et al., 2023). It was first proposed that Prdm9 could have a different role in rattle snake and direct recombination towards CpG islands (Schield et al., 2020), but other observations suggest that the recombination landscapes in these snakes rather reflects the inefficiency of their Prdm9-dependent pathway to direct DSBs away from CpG islands (Hoge et al., 2023). Altogether, the two studies concur on the fact that the recombination landscape in snakes differs from what is observed in mammals, despite the presence of *Prdm9* (Schield et al., 2020; Hoge et al., 2023).

Most of our knowledge about *Prdm9* function and evolution has been acquired in a handful of mammals (mostly human and mice). In mice, Smagulova and colleagues analyzed the position of DSB hotspots in strains carrying different *Prdm9* alleles (Smagulova et al., 2016). Those strains showed differences in their capacity to target DSBs away from the hotspots of the Prdm9^-/-^ mouse (hereafter referred to as ‘MDH’, for ‘Mouse Default Hotspots’). Some strains show a significant deficit of DSB hotspots at MDH loci, while others, carrying less dominant Prdm9 alleles, show up to a 6-fold DSB hotspot enrichment at these loci, even though they represent a small proportion of all recombination hotspots (∼7%) (Smagulova et al., 2016). This shows that even when *Prdm9* is present, Prdm9-independent hotspots can be active in mammals. However, this activity could just be the reflection of a specific Prdm9 deficiency in some mice strains and overall, we still have no clear idea on how prevalent the usage of Prdm9-independent hotspots is in mammals. A comprehensive understanding would require measures of fine-scale variations in recombination rate for a wide range of mammalian species, and ideally across long periods of time.

Recombination hotspots can be mapped directly in meiotic cells (e.g. by chromatin immunoprecipitation with antibodies to DMC1 (Brick et al., 2012; Pratto et al., 2014; Smagulova et al., 2016; Alleva et al., 2021)), but these molecular approaches are tedious and only amenable for a few model organisms. High-resolution recombination maps can also be inferred from patterns of linkage disequilibrium (LD). This approach is more scalable and provides information on sex-averaged historical recombination activity at the population scale. However, this approach remains laborious and expensive (it requires the sequencing of at least 10 individuals per species (Auton and McVean, 2007; Chan et al., 2012)), and it is sensitive to various sources of errors (Spence and Song, 2019; Samuk and Noor, 2022; Raynaud et al., 2023). Hence, for now, such LD-based recombination maps are available only for a very limited number of species.

Alternatively, substitution patterns have been found to be informative about past recombination rates. In particular, it has been shown in mammals that recombination induces a transmission bias of GC alleles through the process of GC-biased gene conversion (gBGC). This eventually leads to an elevation of the WS substitution rate (AT to GC), and a decrease of the SW substitution rate (GC to AT) (Nagylaki, 1983; Duret and Arndt, 2008; Glémin, 2010). The substitution rate matrix can be conveniently summarized by a single parameter, the equilibrium GC-content (hereafter noted *GC*^*\**^), which corresponds to the GC-content that sequences would reach if the pattern of substitution observed in that branch remained constant over time (Duret and Arndt, 2008). *GC*^*\**^ correlates well with the strength of DSB hotspots in mice (Clément and Arndt, 2013), with the LD-based recombination rates in humans (Munch et al., 2014; Glémin et al., 2015), and with the LD-based strength of recombination hotspots in dogs (Axelsson et al., 2012; Auton et al., 2013). Moreover, *GC*^*\**^ reflects the recombination activity along the entire branch where substitution patterns are analyzed, and hence can inform about past recombination events that are no longer detectable with methods measuring recombination in individuals or populations (Lesecque et al., 2014; Munch et al., 2014). This provides insights on the long term use of Prdm9-independent hotspots, integrated over long periods of time, probably encompassing the rise and fall of several *Prdm9* alleles. However, variation in *GC*^*\**^ between species cannot be directly interpreted as variations in recombination rates alone, since *GC*^*\**^ also depends on the mutation bias towards AT, the repair bias towards GC, the effective population size and the mean length of the conversion tracts (Eyre-Walker, 1999; Glémin, 2010).

In this study, we present an estimator of relative recombination rates based on substitution patterns that allows us to directly compare fine-scale recombination rate variations in a wide range of species using only 3 genomes (one focal genome, a sister species and an outgroup). We then use it to assess the recombination activity at MDH loci in 52 species spanning the diversity of boreoeutherians. We reveal a high heterogeneity in the use of these Prdm9-independent hotspots. We show that *Prdm9* alleles in humans and mice have been particularly efficient at directing DSBs away from Prdm9-independent hotspots but that these two species are not representative of all mammals. Finally, we show that three species, namely the southern elephant seal, the ring-tailed lemur and the daurian ground squirrel, have used Prdm9-independent hotspots as much as Prdm9-deficient canids. This shows that the two kinds of hotspots-regulation mechanisms that have been described so far in vertebrates are not mutually exclusive and that the fine-scale recombination landscapes of many mammals are much closer to those of birds and other Prdm9-lacking amniotes than previously thought. We further show that the recombination activity observed at MDH loci in Prdm9-containing mammals depends on the conservation of their DNA methylation pattern, which suggests a link between the evolution of DNA methylation and of Prdm9-independent recombination landscapes.

## Results

### Conservation of recombination hotspots between Prdm9-deficient mammals

In finches and flycatchers, the fine-scale recombination landscape has been shown to be stable through time, as a large proportion of hotspots are shared between closely related species (Singhal et al., 2015; Kawakami et al., 2017). To test whether loci corresponding to Prdm9-independent hotspots are also evolutionary stable in mammals, we analyzed the overlap between recombination hotspots detected in dogs, which naturally lack Prdm9, and those identified in the Prdm9^-/-^ mutant mouse (MDH). Among the 30,929 MDH, 15,009 (49%) could be assigned to one-to-one orthologous loci in the dog genome. Among the 7008 dog hotspots identified in the LD-based recombination map of dogs (Auton et al., 2013), 34% overlap with MDH loci (Suppfig. S1)(compared to 0.06% expected by chance, given that dog hotspots and MDH loci cover respectively 2% and 3% of the dog genome). Although this enrichment is very strong, it should be noted that 42% of the dog hotspots that could be mapped on the mouse genome occur outside of MDH loci (1809/3109) (Suppfig. S1). This number is difficult to interpret because it has been shown that LD-based methods can produce a high number of false positives (Raynaud et al., 2023). Moreover, DSB hotspots are obtained on males only, while LD maps are sex-averaged. It is also possible that some MDH loci have not been identified as hotspots in dogs simply because they did not meet the threshold criteria to be defined as such. To avoid the problem of the arbitrary threshold, we computed the LD-based recombination rate in dogs (from Auton et al. (2013)) as a function of the distance to the closest MDH loci (Fig. 1A&B) (Auton et al., 2013). We divided the 30,929 mouse Prdm9^-/-^ hotspots into three equally sized categories of strength, based on DMC1 ChIP-seq read counts (Smagulova et al., 2016). Respectively 5,266 strong MDH, 4,961 medium MDH and 4,781 weak MDH could be mapped on the dog genome. There is a sharp peak of recombination centered on MDH loci in dogs. This peak is higher for strong MDH and weaker for medium and weak ones (Fig. 1A). This confirms that many recombination hotspots are conserved between Prdm9^-/-^ mice and dogs, but it does not rule out the existence of species-specific recombination hotspots. To explore factors that might drive the evolution of Prdm9-independent hotspots, we analyzed their DNA methylation level in the germline. Indeed, indirect evidences suggested that recombination hotspots in dogs are associated to germline hypomethylated regions (HMRs) (Berglund et al., 2015). Using HMRs identified by bi-sulfite sequencing in dog sperm (Qu et al., 2018), we observed that 74% of dog hotspots are located inside HMRs, which represent only 3.7% of the dog genome. In mice, the overlap is even stronger: using HMRs identified in mouse sperm (Hammoud et al., 2014), we observed that out of the 30,929 hotspots found in the Prdm9^-/-^ mutant, 93% are located within HMRs, which cover only 4.6% of the mouse genome (see methods for details). This indicates that Prdm9-independent hotspots are associated with DNA hypomethylation both in Prdm9^-/-^ mutant mice and in canids. Interestingly, 48% of MDH loci are methylated in dog sperm if we restrict the definition of hotspots to their midpoint (7,186/15,009). This shows that many MDH loci are specifically hypomethylated in mice but not in dogs. This is consistent with previous observations showing that murid genomes have accumulated many new HMRs compared to other mammals (Qu et al., 2018). To test whether these shifts in methylation levels are associated with changes in recombination activity, we computed the LD-based recombination rate in dogs as a function of the distance to the closest MDH locus, separating those whose midpoints overlap a HMR in dogs (7,186), and those that do not (7,283)(Fig. 1B). There is a high and pronounced recombination peak at MDH loci that are hypomethylated in dog sperm (Fig. 1B). In contrast there is almost no elevation of recombination at MDH loci that are methylated in dogs (Fig. 1B). This confirms that methylation is clearly associated to recombination hotspots in the absence of Prdm9, and that many Prdm9-independent recombination hotspots are species-specific.

**Figure 1:**
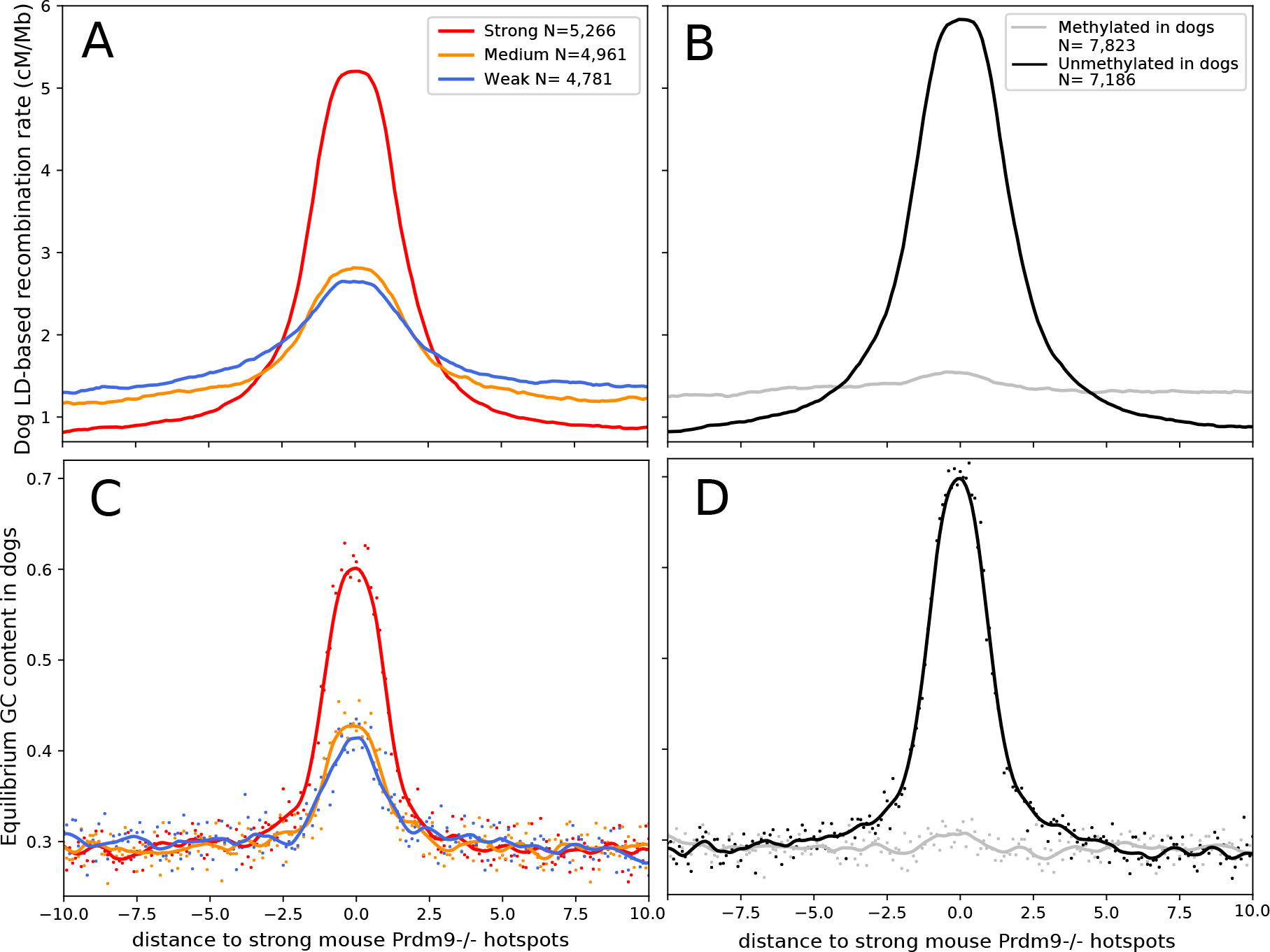
A) Dog LD-based recombination rate as a function of the distance to the closest MDH loci. MDH were divided in three equally sized categories of strength: strong hotspots in red (10-190 FPKM), medium hotspots in orange (5-10 FPKM) and weak hotspots in blue (0-5 FPKM). The line directly correspond to the mean value of LD-based recombination rate in a 100 bp window. B) Same as A but MDH loci were divided in two categories depending on their methylation level in dog sperm. MDH loci that are found outside hypomethylated regions in dogs are in grey and those found inside hypomethylated regions are in black. C) & D) Equilibrium GC content in dogs as a function of the distance to the closest MDH loci. Using the same partitions of hotspots as A) and B). Points correspond to the mean value of *GC*^*\**^ in a 100 bp window. The line correspond to a smoothing of the data with a loess function.

### Equilibrium GC content is a good predictor of past recombination activity

To assess whether equilibrium GC content could be used as a proxy of recombination rate in other species, we repeated the above analyses using *GC*^*\**^ instead of LD-based recombination rate (Fig. 1C&D). Strikingly, the profile of *GC*^*\**^ perfectly mirrors the LD-based recombination rate profile. It should however be noticed that the *GC*^*\**^ peaks are slightly sharper than the LD-based ones (Fig. 1C&D), as are the peaks of DMC1 ChIP-seq read coverage in the Prdm9^-/-^ mouse (Suppfig. S2). This suggests that *GC*^*\**^ is able to capture signals of past recombination with a higher spatial resolution than LD.

### Estimation of the relative recombination rate in Prdm9-independent hotspots

Following a large body of literature, we showed that *GC*^*\**^ can be very informative on intra-genomic recombination rate variations (Pessia et al., 2012; Auton et al., 2013; Clément and Arndt, 2013; Lartillot, 2013; Munch et al., 2014; Glémin et al., 2015; Singhal et al., 2015; Figuet et al., 2015; Bolívar et al., 2016; Galtier et al., 2018; Charlesworth et al., 2020). However, the height of the *GC*^*\**^ peak in hotspots is difficult to interpret in term of recombination rate because it is also affected by other parameters (the length of gene conversion tracts, the mutation bias towards AT, the mismatch repair bias towards GC and the effective population size), which can vary between species (Lartillot, 2013; Galtier et al., 2018; Galtier, 2021). We thus derived an estimator that can capture the relative recombination rate at Prdm9-independent hotspots that controls for those parameters and is therefore comparable between species. Using the probability of fixation of AT and GC alleles in presence of gBGC derived by Nagylaki (1983), we obtained an expression of the ratio of the recombination rate within hotspots relative to their flanking regions (see details in the Methods). This relative recombination rate only depends on *GC*^*\**^ inside hotspots and in flanking regions, and on the mutation bias (*GC*^*μ*^).

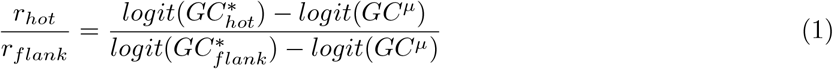

where *r*_*hot*_ is the recombination rate in hotspots, *r*_*flank*_ the recombination rate in flanking regions and *GC*^*μ*^ the GC content expected under mutation only. It should be noted that *r*_*hot*_ and *r*_*flank*_recombination events that can lead to gBGC (potentially COs and/or NCOs).

For the rest of the study, hotspots are defined as the 400 bp regions centered on their midpoint, and flanking regions as those spanning from 5 to 8 kb upstream and downstream of the hotspots (Fig. 2A).

**Figure 2:**
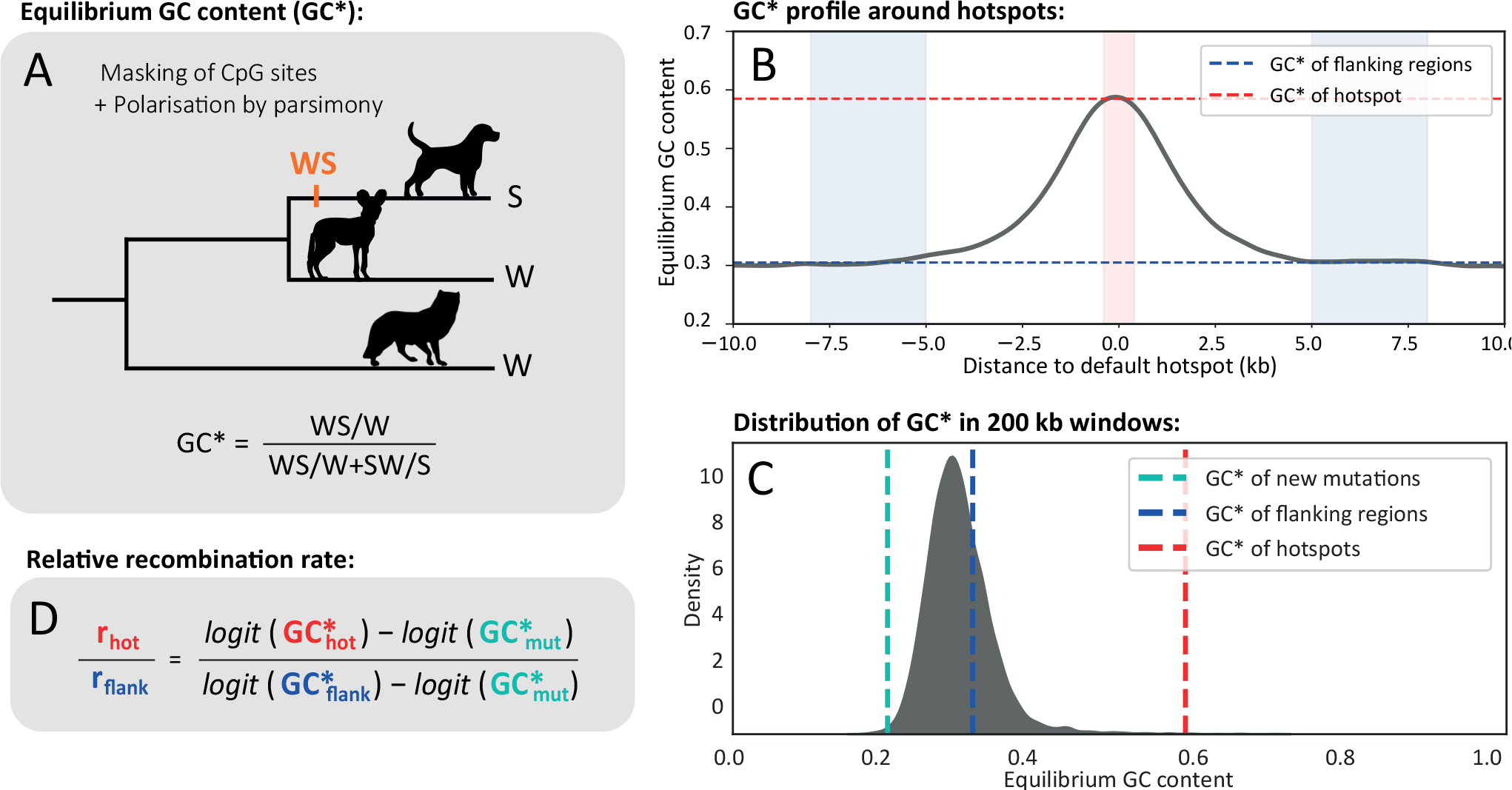
Overview of the method for inferring relative recombination rates in Prdm9-independent hotspots. A) We call substitutions using parsimony on trios of closely related species after having masked CpG dinucleotides. B) We compute *GC*^*\**^ in 400 bp windows centered on the midpoint of the Prdm9-independent hotspots, and *GC*^*\**^ in the flanking regions (from 5 to 8 kb upstream and downstream of the center of the Prdm9-independent hotspot. C) We compute the distribution of *GC*^*\**^ in 200 kb windows and take the 1^*st*^ percentile as the *GC*^*\**^ of new mutations. D) Using the probability of fixation of AT and GC alleles in presence of gBGC derived by Nagylaki (1983), we compute the relative recombination rate as a function of the three values of *GC*^*\**^ (see methods).

Using the three values 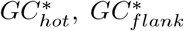 and *GC*^*μ*^, it is possible to compute a measure of the relative recombination rate within hotspots compared to their flanking regions, which can then be compared between species. Equation 1 holds true under the assumption that the other four parameters affecting the strength of gBGC (effective population size, mismatch repair bias, length of the conversion tract and the mutation bias) do not differ between the hotspots and their flanking regions (see Methods). As shown in the previous section, Prdm9-independent hotspots are often hypomethylated. This implies that the mutation rate from CpG to TpG or CpA in hotspots is lower than in the flanking regions, which violates our assumption of a constant mutation bias. To avoid this problem, we excluded CpG sites from all the analyses.

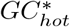 and 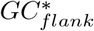 can easily be computed from the substitutions in Prdm9-independent hotspots and their flanking region but *GC*^*μ*^ is more difficult to estimate. Interestingly, it has been shown that genome-wide variations of *GC*^*\**^ are mainly the result of gBGC, and that *GC*^*μ*^ is quite constant along chromosomes, despite large-scale variation in mutation rates (Smith et al., 2018). An approach to estimate *GC*^*μ*^ consists in measuring variation in substitution patterns along the genome, and to consider the regions of the genome with the lowest *GC*^*\**^ as a proxy for *GC*^*μ*^ (Lartillot, 2013). Following this logic, we divided the genome of each species in windows of 200kb, and defined *GC*^*μ*^ as the value of the first percentile of the distribution of *GC*^*\**^ to avoid outliers (Fig. 2B) (see methods for detailed justifications). This method allows us to estimate *GC*^*μ*^ for a wide range of species, simply based on substitution patterns in the terminal branch.

### Prdm9-independent recombination hotspots are active in most mammals

Using this estimator of relative recombination rate, we assessed whether MDH loci showed an enrichment of recombination in other mammals. We identified MDH orthologous loci in the genome of 51 other mammals and estimated the relative recombination rates at these loci using the method described above (gBGC-based relative recombination rates). Around 75% of the species (39/52) show a significant enrichment of recombination in MDH loci compared to flanking regions (Fig. 3). In 77% of those species (30/39), the recombination activity at MDH loci is conserved, with strong MDH loci showing a significantly higher recombination enrichment than weak ones (Fig. 3). The remaining 23% of species (9/39) show a lower recombination activity in Prdm9-independent hotspots. Therefore, we might be lacking statistical power to confirm a conservation of hotspot strength in those species (Fig. 3). Interestingly, in 10% of species (5/52), including mice, there is less recombination in strong Prdm9-independent hotspots compared to weak ones, which is consistent with the active deviation of recombination away from those sites observed in those mice having the most dominant Prdm9 alleles (Smagulova et al., 2016). The causes for this active deviation are however still not clear (Smagulova et al., 2016).

**Figure 3:**
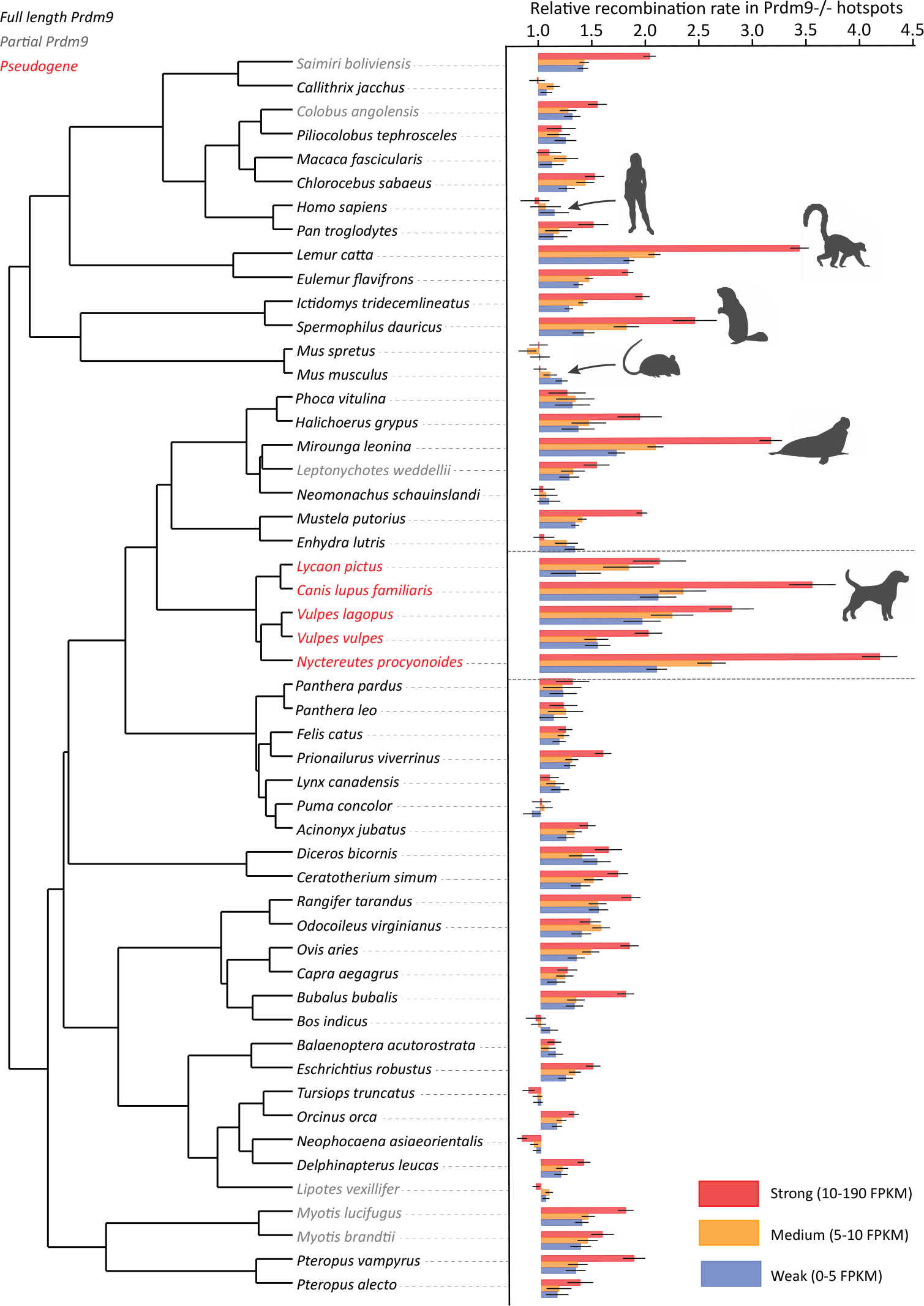
Relative recombination rates at at loci orthologous to mouse Prdm9^-/-^ DSB hotspots (MDH loci) in 52 mammals. MDH loci were binned in 3 equally sized categories of strength based on the number of DMC1 Chip-seq reads of Prdm9^-/-^ DSB hotspots. The number of MDH loci for each category varied from ∼4,000 in *Myotis brandtii* to ∼9,000 in *Mus spretus* (see details in Supplementary Table 1). The tree has been retrieved from TimeTree5 (Kumar et al., 2022). Species with a complete Prdm9 are written in black, in grey species for which we failed to find a complete Prdm9 in the reference genome assembly, and in red the 5 canids (where Prdm9 is a pseudogene). Error bars correspond to a 95% confidence interval obtained by bootstrapping the substitutions for computing 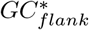 and 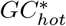. We considered the recombination enrichment to be significant if the confidence interval is above one.

To test whether as in dogs, methylation plays a role in determining hotspots in other mammals, we separated MDH loci in two subsets: loci for which the hypomethylation pattern is conserved between mice and dogs (and thus likely to be consistently hypomethylated in other mammals as shown in Qu et al. (2018)), and loci that are hypomethylated in mouse but not dogs (a majority of which corresponding to mouse-specific HMRs (Qu et al., 2018)). For MDH loci whose hypomethylation pattern is not conserved, we observed a very weak increase in recombination in all species (Suppfig. S4).

Conversely, MDH loci with a conserved hypomethylation pattern, show contrasting levels of recombination activity: 25% of species (13/52, including mice and humans) show a deficit of recombination at these loci, 12% (6/52) show no elevation of recombination, whereas 63% (33/52) show a strong recombination activity. This indicates that DNA hypomethylation is associated with a deficit of recombination in some species (such as mice and humans), while it is associated with recombination hotspot in others. This latter group includes the 5 canids (which all lack Prdm9), but also many other mammals with an intact Prdm9. This shows that DNA hypomethylation can be associated with recombination hotspots even in the presence of Prdm9.

To get further insight into the evolution of Prdm9-independent hotspots in mammals, we measured the relative activity at loci orthologous to dog LD-based recombination hotspots (DRH for ‘Dog Recombination Hotspots’) in the 51 other mammals. We observed a strong correlation between the recombination activity at MDH loci and DRH loci (Suppfig S5), which is expected since they largely overlap. Nevertheless, species that are phylogenetically closer to dogs show higher recombination activity in DRH loci whereas those phylogenetically closer to mouse show higher recombination activity in MDH loci (Suppfig S5). This confirms that despite a general conservation, Prdm9-independent hotspots are still evolving in mammals.

Canids, which all lack Prdm9, dominate the list of species that exhibit the highest recombination levels at MDH loci, holding top ranks out of 52 (Fig. 4A). This finding both shows that the recombination landscape is stable in canids as it is in passerines, and validates our relative recombination rate estimator (Fig. 4A). Interestingly, there are several other mammals that show a similar enrichment of recombination activity at MDH loci. Notably, three species show a recombination activity at MDH loci significantly higher than some of the canids: ring-tailed lemurs (*Lemur catta*), southern elephant seals (*Mirounga leonina*) and daurian ground squirrels (*Spermophilus dauricus*) (Fig. 4A).

**Figure 4:**
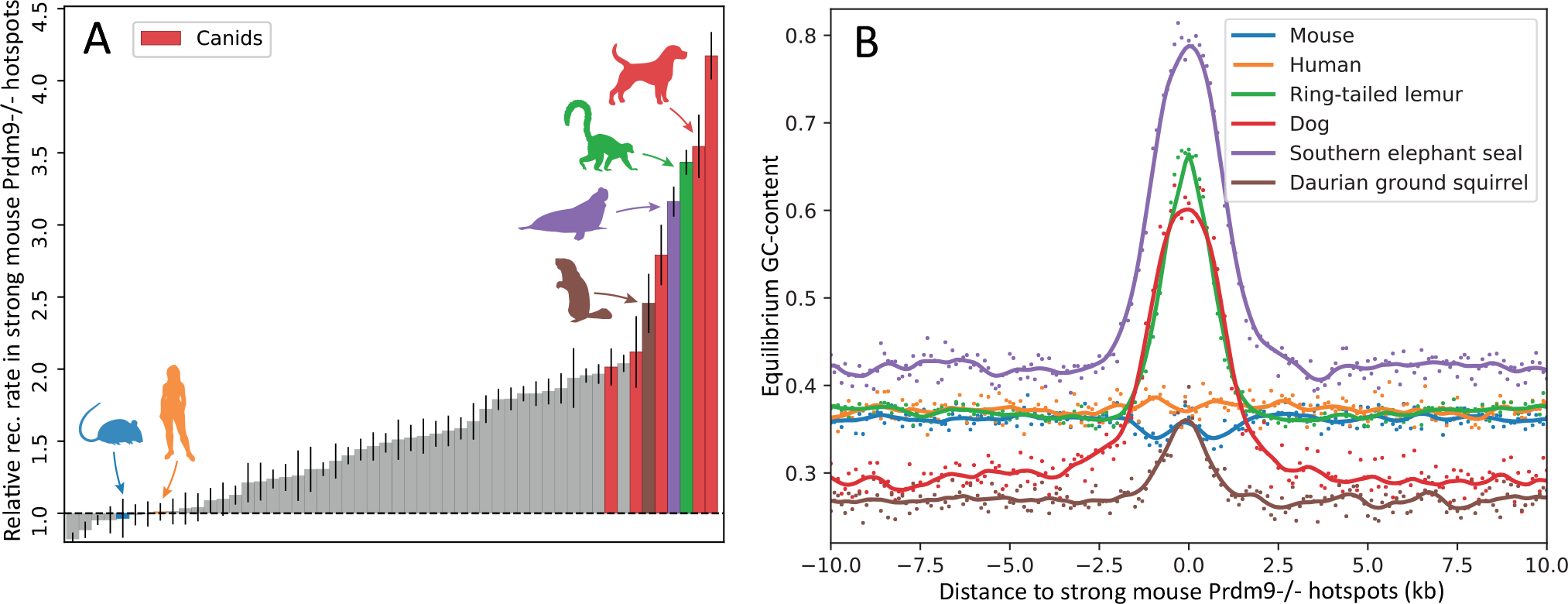
(A): Sorted relative recombination rate in strong DSB hotspots (*>*10 FPKM) of a Prdm9 -/- mouse in 52 mammals. Error bars correspond to a 95% confidence interval obtained by bootstrapping the substitutions for computing 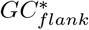 and 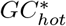. (B): *GC*^*\**^ as a function of the distance to the center of the closest MDH locus for humans, mice, dogs, and the three outlier species. Each point corresponds to an estimation of *GC*^*\**^ in a 100 bp window.

It should be noted that these three species encode a full-length Prdm9, encompassing the four protein domains (KRAB, SSXRD, SET and the zinc finger array). The Prdm9 allele represented in the reference genome assembly contains 11 zinc fingers in daurian ground squirrels, and 14 zinc fingers in ring-tailed lemurs. For the southern elephant seal, there is only one complete zinc finger represented in the reference genome because of an assembly gap within the array. In ring-tailed lemurs and daurian ground squirrels, the comparison of zinc finger sequences showed an excess of amino-acid changes relative to the neutral expectation, particularly at positions -1, 3, and 6 that are involved in DNA binding, (SuppFig. S6).

This suggests that Prdm9 has been subject to positive selection in these two lineages, which is suggestive of the red-queen dynamic that is expected when Prdm9 determines the location of recombination hotspots. Thus, Prdm9 does not show any sign of pseudogenization or functional change in these three species where Prdm9-independent hotspots appear to have been particularly active.

As an illustration of our substitution-based approach, we plotted *GC*^*\**^ as a function of the distance to the closest strong MDH loci in dogs, humans, mice and the three outlier species (Fig. 4B). We observed no elevation of *GC*^*\**^ in humans and mice in strong MDH loci, confirming that very little recombination occurred in their lineage (Fig. 4B). In dogs, ring-tailed lemurs, southern elephant seals and daurian ground squirrels, there is a pronounced peak of *GC*^*\**^ at strong MDH loci (Fig. 4B).

## Discussion

The current paradigm is that vertebrates either possess a full-length functional Prdm9, recombine away from promoter-like features, and display a fast evolving recombination landscape, or they lack a functional Prdm9 in which case they consistently recombine in CpG islands (e.g. in canids, birds or the swordtail fish) (Baker et al., 2017). Our results revealed that there is a continuum between these two types of recombination landscapes and that despite the presence of Prdm9, some species use Prdm9-independent hotspots as much as canids. We showed that the activity of those Prdm9-independent hotspots is highly dependent on DNA hypomethylation. This implies that despite a general conservation, Prdm9-independent recombination hotspots are evolving slowly, in concert with germline DNA hypomethylation (Qu et al., 2018; Berglund et al., 2015).

From a methodological perspective, while signatures of gBGC have been commonly used in studies of recombination landscapes (Axelsson et al., 2012; Auton et al., 2013; Munch et al., 2014; Lesecque et al., 2014; Singhal et al., 2015; Charlesworth et al., 2020; Hoge et al., 2023), the approach presented here allows to quantify gBGC-based relative recombination rates along a branch that are comparable between species. The analysis of gBGC signatures only requires the genome of three closely related species, and is able to detect recombination activity at a given set of loci with a very high spatial resolution, even better than LD-based methods (compare Fig. 2D and 2B). It thus offers the possibility for large-scale comparative studies of fine-scale recombination landscapes. However, this approach requires a large number of substitutions to estimate *GC*^*\**^ precisely and is therefore not appropriate to measure recombination at a single locus.

Moreover, it should be noted that our estimation of recombination activity using gBGC does not allow one to conclude on the nature of recombination events (COs or NCOs) that we detect at Prdm9-independent hotspots. In humans, there is evidence that both COs and NCOs induce gBGC but in mice, only NCOs appear to do so (Williams et al., 2015; Arbeithuber et al., 2015; Halldorsson et al., 2016; Li et al., 2019). This suggests that the type of recombination events triggering gBGC can vary among mammals. Thus, it is possible that while a large number of DSB occur at Prdm9-independent hotspots in our three outlier species, COs still tend to be associated to Prdm9-directed hotspots. Altogether, while the enrichment of recombination events in Prdm9-independent hotspots is very clear in numerous mammals, the way they are repaired remains to be explored.

### On the maintenance of a default hotspot regulation mechanism

It had been previously demonstrated that both Prdm9-dependent and Prdm9-independent pathways coexist in two snakes genera (Hoge et al., 2023). The authors suggested that a change in the binding affinity of a gene which operates downstream of Prdm9 could explain this coexistence and that selection might be operating on these genes to fine-tune the usage of Prdm9-independent hotspots in vertebrates (Hoge et al., 2023).

In placental mammals we revealed that the coexistence of both Prdm9-dependent and Prdm9-independent pathways to direct DSBs is pervasive. We showed that those pathways determine recombination hotspots with varying proportions across species. Interestingly, these variations do not show a strong phylogenetic structure, suggesting that this evolution can be very rapid. Furthermore, the species with the highest levels of Prdm9-independent hotspot usage have quite contrasted life history traits, which they mostly share with sister species with lower Prdm9-independent hotspot usage. Thus, it is difficult to imagine that differences of Prdm9-independent usage depend on reproductive life history traits. Moreover, in mice, even if the usage of Prdm9-independent hotspots is quite low overall, there exist substantial variations that seem to depend only on Prdm9 alleles (Smagulova et al., 2016). A plausible explanation for the maintenance of the Prdm9-independent pathway in placental mammals would be that variation in Prdm9-independent hotspots usage depends on the efficacy of Prdm9 alleles to recruit the DSB machinery.

Overall, proper chromosome pairing can be achieved through the two different pathways mentioned above (Prdm9-dependent or Prdm9-independent). When the efficiency of one pathway is altered, better chromosome pairing can be restored either by a mutation restoring the efficiency of the altered pathway, or by a mutation increasing the efficiency of the other pathway. For the Prdm9 pathway, we know that alleles inevitably decrease in efficiency due to the erosion of their high affinity targets, which reduces the probability of symmetrical binding, and thus impairs efficient chromosome pairing (Baker et al., 2015, 2022; Latrille et al., 2017; Genestier et al., 2023). This efficiency can be restored either by a new Prdm9 allele inducing a red-queen dynamic (Úbeda and Wilkins, 2011; Latrille et al., 2017; Baker et al., 2022; Genestier et al., 2023), but also by a mutation increasing the efficiency of the Prdm9-independent pathway. Every mutation increasing the efficiency of the Prdm9-independent pathway lessens the deleterious effect of a mutation that reduces Prdm9 efficiency. Conversely, new efficient Prdm9 alleles will lessen the deleterious effect of a mutation that decreases the efficiency of the Prdm9-independent pathway.

Of note, if this dynamic reaches a point where the Prdm9-independent pathway becomes sufficient for correct chromosome pairing, *Prdm9* can be lost without strong fitness consequences. Under this model, *Prdm9* is lost through the accumulation of small effect mutations which reduce its utility, rather than a sudden loss that would imply very inefficient selection. The continuum in the use of Prdm9-independent hotspots we observe in mammals could reflect different stages along this path, and despite still having a fully functional *Prdm9*, the three outlier species could be on their way of losing it. It is also possible that their *Prdm9* have only been going through a temporary inefficient phase compensated by the Prdm9-independent pathway, but has now been rescued by a new efficient Prdm9 allele.

### The recombination landscape of amniotes

Overall, our results suggest that in addition to Prdm9-directed hotspots, many mammals share some of their recombination hotspots with other amniotes, (Axelsson et al., 2012; Singhal et al., 2015; Schield et al., 2020; Hoge et al., 2023), and therefore the fine-scale recombination landscapes of mammals, birds and snakes is probably more similar than previously thought (Baker et al., 2017). However, the determinants of Prdm9-independent hotspots usage remain unclear. Interestingly DNA methylation has also been found to be a suppressor of recombination in a fungi hotspot (Maloisel and Rossignol, 1998), in plants (He et al., 2017; Choi et al., 2018), and in honey bees (Wallberg et al., 2015), which suggests that local hypomethylation is a common determinant of recombination hotspots in eukaryotes. However, the association between hypomethylation and recombination has not been formally established in non-mammalian amniotes and remains to be tested. Moreover, this association need not be causal, as the potentially diverse molecular mechanisms of Prdm9-independent recombination in amniotes remain largely unknown. In particular, the results presented here suggest that despite having lost Prdm9, red foxes (*Vulpes vulpes*) and african wild dogs (*Lycaon pictus*) have only a mild recombination enrichment in Prdm9-independent hotspots compared to other canids. It has been previously noted that the number of recombination hotspots varies between Prdm9-deficient amniotes. Notably in finches and flycatchers, only few LD-based hotspots have been reported compared to dogs (Auton et al., 2013; Singhal et al., 2015; Kawakami et al., 2017). Altogether, it is still not clear what drives the concentration of recombination events in absence of Prdm9, and why some species have numerous hotspots and others less.

Finally, the widespread use of Prdm9-independent recombination hotspots demonstrated in the present study is likely to have important consequences for genome evolution. In particular, the fact that gBGC is stronger in hypomethylated regions in numerous mammals and in several passerines provides a convincing explanation for the widespread GC-richness of CpG islands in amniotes.

## Material & Methods

### Prdm9-independent hotspots and DNA methylation datasets

Data on the location of DSB hotspots in Prdm9^-/-^ knock-out mice, detected by DMC1 ChIP-seq experiment, were retrieved from the study by Smagulova et al. (2016). The dog LD-based recombination map and the position of hotspots were retrieved from the study of Auton et al. (2013). We excluded hotspots that were larger than 20 kb as they do not fit the definition of hotspots. Data sets of hypomethylated regions in mouse and dog sperm (identified using Bi-sulfite sequencing) were retrieved from the literature (Hammoud et al., 2014; Qu et al., 2018). Even though methylation data on spermatocytes would have been more fit for the task at hand, only sperm is available in the literature for dogs.

### Whole genome alignments

For most mammals except canids, felids and phocids, whole genome alignments (WGAs) were obtained from Genereux et al. (2020). In order to get further phylogenetic resolution in canids, and in closely related outgroups, we generated a WGA of high quality genomes for 19 carnivores downloaded from NCBI (Supplementary Table 3), using the Progressive Cactus aligner (v1.3.0) (Armstrong et al., 2020). We first defined a “guide” species tree using the topology obtained from TimeTree5 (Kumar et al., 2022). To streamline the computational process, we ran Progressive Cactus separately for canids, felids and phocids species, using different “–root” options on the same guide tree. We created a root alignment by running Progressive Cactus with the inferred ancestral genome of each of the three clades. We obtained the final WGA using the “halAppendSubtree” command to iteratively include the three sub-alignments at the corresponding ancestral nodes (Hickey et al., 2013).

### Defining orthologous regions

To find the orthologous regions of the Prdm9-independent hotspots in the genomes of other mammals we used halLiftover (Hickey et al., 2013). We first made a liftover from the mouse/dog genome to the target genome using the midpoint of each feature and removed multi-mapping features. Then we lifted back the single-mapping features from the target genome to the dog/mouse genome and again removed multi-mapping features. This approach ensures that all orthologous loci were one-to-one.

### Hotspots overlap

We considered hotspots to be overlapping if their midpoint was at less than 5 kb one from another. This is equivalent to a strict overlap for hotspots defined as 5 kb windows centered on their midpoint. Using this approach, we calculated the percentage of the genome covered by the hotspots by multiplying the number of hotspots by 5,000, and dividing by the assembly size. For the overlap with HMRs we defined hotspots as the 5 kb windows centered on their midpoint and kept the size of HMRs defined in the study of Qu et al. 2018 (Qu et al., 2018). The percentage of the genome covered by HMRs was computed as the sum of all HMR sizes divided by the assembly size.

### Substitution mapping

We selected trios of closely related species such that the divergence between the three species was low enough to avoid double substitutions, but with an outgroup distant enough to avoid incomplete lineage sorting based on the guide tree used in the Zoonomia WGA (Genereux et al., 2020). We tried to take trios spanning the diversity of boreoeutherians, avoiding over-sampling of disproportionately represented groups (primates and artiodactyles). A complete list of the trios used are available in Supplementary table 4. *A posteriori*, it appeared that there were substantial variations between the branch lengths used to map substitutions, but we showed that the divergence was still low enough (*<* 2.5%) and did not influence our result (Suppfig. S7). Genome quality was very variable. We thus controlled that genome quality (approximated by the N50 statistics) did not influence our results (Suppfig. S8). To call substitutions, we retrieved multispecies alignment using hal2maf (Hickey et al., 2013). We excluded alignment blocks which size was inferior to 50 bp to avoid poorly aligned regions, and duplicated regions. We then excluded CpG sites (sites for which at least one of the three species has a CpG) to avoid convergent mutations. Finally, we called substitutions using parsimony as depicted in Fig. 2.

### Measures of equilibrium GC content

We compute the equilibrium GC content as follows:

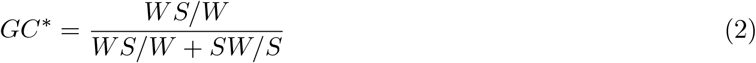

With *W* the AT content of the region (CpG masked), *S* the GC content of the region (CpG masked), *WS* the number of Weak to Strong substitutions and *SW* the number of Strong to Weak substitutions.

### LD-based recombination rate and GC^*\**^ profiles around hotspots

We cut the genome in windows of 100 bp. We extracted the LD-based recombination rate for each window. We also computed the distance between the midpoint of the window and the closest midpoint of a hotspot using bedtools closest (Quinlan and Hall, 2010). We then made bins of distances to the closest hotspot every 100 bp. For *GC*^*\**^, we repeated the same procedure, but we computed the counts of WS and SW substitution for each window, and then computed *GC*^*\**^ using the total substitutions count for all windows in each distance bin as described in the previous section.

### Estimation of the mutation bias

Germline mutations rates have been measured by sequencing parent–offspring trios in 36 mammalian species (Bergeron et al., 2023). However, for most species, the number of detected mutations is too limited (typically less than 100 de novo mutations) to estimate *GC*^*μ*^ accurately. Thus, measures of *GC*^*μ*^ based on empirical data are associated with very large confidence intervals (Wong et al., 2016; Milholland et al., 2017; Wang et al., 2020). Even when there is enough statistical power, the results vary substantially between different datasets (Wong et al., 2016; Milholland et al., 2017). In addition to the issue of reproducibility, it has been demonstrated that the mutation spectra can vary rapidly within human populations (Harris and Pritchard, 2017). Therefore, *GC*^*μ*^ estimated from living individuals may not necessarily reflect the average *GC*^*μ*^ in the terminal branch. We therefore took a similar approach to (Lartillot, 2013). We divided the genomes in windows of 200 kb and took the value of the first percentile of *GC*^*\**^ as an estimate of *GC*^*μ*^. To ensure that our results were not sensitive to our estimation of *GC*^*μ*^, we used different thresholds to compute it (Suppfig. S3B&C), and recovered the same results. We also controlled that our gBGC-based relative recombination rates were not correlated to our estimations of *GC*^*μ*^ (Suppfig. S3A).

### Estimation of relative recombination rates from GC^*\**^

Using a Wright-Fischer diffusion approximation and assuming that mutations are selectively neutral, the rate of Weak-to-Strong substitution in a given branch can be written as follows (Nagylaki, 1983)

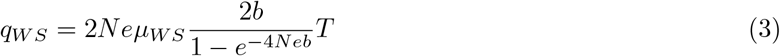

where *μ*_*SW*_ is the mutation rate per generation from W to S, *b* the gBGC coefficient, *T* the divergence time from the ancestral node in generations, and *Ne* the effective population size.

The gBGC coefficient is directly linked to the recombination rate, with *b* = *b*_0_*rl* where *r* is the recombination rate per base pair per meiosis, *b*_0_ the repair bias, and *l* the length of the conversion tract in base pair. It should be noted that *r* encompasses all recombination events that can lead to gBGC (CO and/or NCO)

Similarly, the rate of Strong-to-Weak substitutions can be written as follows:

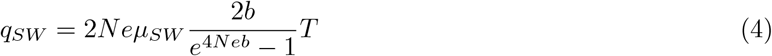

The equilibrium GC content can be written as follows:

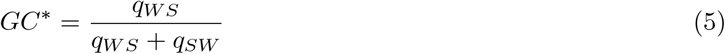

Thus, we can write:

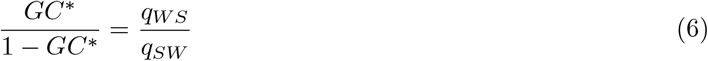

Simplifying the previous equations we obtain:

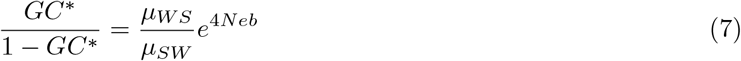

Thus, the population-scaled gBGC coefficient. (*B* = 4*N eb*) can be written as:

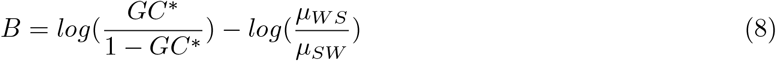

or:

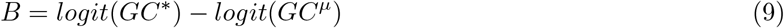

where *GC*^*μ*^ is the equilibrium GC content under the mutational bias only 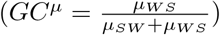.

Let us note *B*_*hot*_ the population-scaled gBGC coefficient within hotspots:

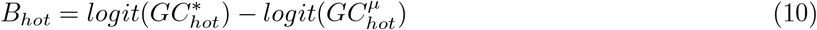

And *B*_*flank*_ the population-scaled gBGC coefficient in their flanking regions (defined here as 3kb-long segments, located at 5-kb of the hotspot center, Fig. 2A)

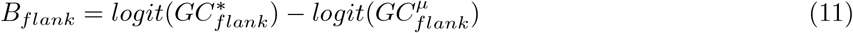

*B* depends on *b*_0_, *N*_*e*_, *l*and *r* (*B* = 4*N erb*_0_*l*). The first three parameters (*b*_0_, *N*_*e*_, *l*) are not expected to differ between the hotspot and their flanking regions. Thus, the ratio between the recombination rate within hotspot (*r*_*hot*_) over the recombination rate in their flanking regions (*r*_*flank*)_can be written as:

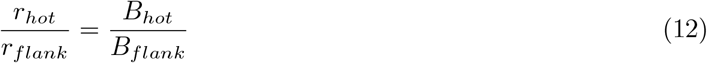

And thus, under the assumption that the mutational bias does not differ between hotposts and their flanking regions (i.e. 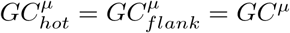)

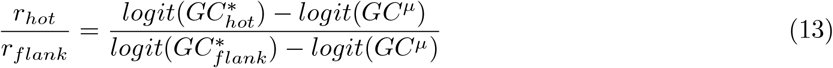

### Annotation of Prdm9 in mammals

We investigated the presence of Prdm9 homologs in each of the 52 species analyzed. The full-length Prdm9 isoform encompasses four domains (KRAB, SSXRD, SET and the zinc finger array). It is encoded by 10 exons (corresponding to exons 2 to 11 of human Prdm9, exon 1 being within the 5’UTR): exons 3 and 4 encode the KRAB domain, exon 7 encodes the SSXRD domain, exons 8-10 encode the SET domain, and exon 11 encodes the zinc finger array. We first searched for Prdm9 homologs by sequence similarity (Camacho et al., 2009) against mammalian proteins annotated in RefSeq (https://www.ncbi.nlm.nih.gov/blast/), using the human protein (NP 001363829.1) as a query. We performed a multiple alignment of the strongest hits, to assess their completeness: homologs where considered as complete if they encompassed the 10 protein-coding exons, from the start codon, up to the beginning of the zinc finger array. By this approach, we identified complete Prdm9 homologs in 16 species (Supplementary Table 2). For the 36 other species, we further analyzed the corresponding reference genome to identify potential Prdm9 homologs. We performed a TBLASTN search against reference genomes, using the 16 previously identified Prdm9 as queries, and extracted loci presenting hits with the zinc finger domain and with at least 2 other Prdm9 exons, within less than 100 kb. Then, for each candidate genomic fragment, we used GeneWise (Birney et al., 2004) to annotate protein-coding regions by similarity with a representative complete Prdm9 protein taken from a closely related species. Of note, GeneWise does take frameshifts into account, and is therefore appropriate to annotate both genes and pseudogenes. In most species we identified one single candidate locus per genome. When several loci were found, we retained the one(s) encoding the most complete protein. By this approach, we identified Prdm9 loci in all 36 species, 25 of which encode a complete Prdm9 protein. Thus, in total, we identified complete Prdm9 proteins in 41 of the 52 species analyzed. In agreement with previous reports ((Axelsson et al., 2012; Auton et al., 2013)), we found Prdm9 to be pseudogenized in the 5 canids (*Lycaon pictus, Canis lupus familiaris, Vulpes lagopus, Vulpes vulpes, Nyctereutes procyonoides*). In the 6 remaining cases, we failed to annotate a complete Prdm9 protein: either several exons were missing (*Myotis lucifugus, Myotis brandtii, Leptonychotes weddellii, Colobus angolensis*), or the gene contained one exon with a frameshifting mutation (respectively in exon 8 in *Saimiri boliviensis* and in exon 10 in *Lipotes vexillifer*). In absence of data from more individuals, it is difficult to state whether these cases result from sequencing errors or assembly artefacts or if they correspond to bona fide pseudogenes. We therefore tentatively annotate these 6 cases as ‘partial’ Prdm9. The number of zinc fingers in the annotated proteins vary from 0 to 14. This number should however be considered with caution because the zinc finger array is encoded by a highly polymorphic minisatellite repeat, which is prone to errors during genome assembly. The detailed list of Prdm9 sequences is given in Supplementary Table 2 and the corresponding protein multiple alignment is given in Supplementary Data 1.

## Acknowledgments

We wish to thank Nicolas Lartillot, Carina Farah Mugal, and Bernard de Massy for very useful reviews on a previous version of this manuscript, and Anaïs Duhamel for help with the figures. This work was performed using the computing facilities of the CC LBBE/PRABI.

## Author contributions

Original idea: J.J.; Model conception: J.J.; Code: J.J., D.P, A.L, T.T and L.D; Data analyses: J.J., D.P, A.L, T.T and L.D; Interpretation: J.J., D.P, A.L, T.T and L.D; First draft: J.J.; Editing and revisions: J.J., D.P, A.L, T.T and L.D; Funding: L.D.

## Data availability

Analysis scripts and documentation for whole genome alignments are available at https://github.com/AlexandreLaverre/CANCGI-WGA/ The hal file of the Carnivore WGA and analysis scripts and documentation for Prdm9 annotation and study of positive selection are available at https://zenodo.org/records/10149667. The rest of analysis scripts and documentation are available at https://gitlab.in2p3.fr/julien.joseph/defhot.

## References

Alleva, B., Brick, K., Pratto, F., Huang, M., and Camerini-Otero, R. D. (2021). Cataloging Human PRDM9 Allelic Variation Using Long-Read Sequencing Reveals PRDM9 Population Specificity and Two Distinct Groupings of Related Alleles. Frontiers in Cell and Developmental Biology, 9.

Arbeithuber, B., Betancourt, A. J., Ebner, T., and Tiemann-Boege, I. (2015). Crossovers are associated with mutation and biased gene conversion at recombination hotspots. Proceedings of the National Academy of Sciences, 112(7):2109–2114.

Armstrong, J., Hickey, G., Diekhans, M., Fiddes, I. T., Novak, A. M., Deran, A., Fang, Q., Xie, D., Feng, S., Stiller, J., Genereux, D., Johnson, J., Marinescu, V. D., Alföldi, J., Harris, R. S., Lindblad-Toh, K., Haussler, D., Karlsson, E., Jarvis, E. D., Zhang, G., and Paten, B. (2020). Progressive Cactus is a multiple-genome aligner for the thousand-genome era. Nature, 587(7833):246–251.

Auton, A., Fledel-Alon, A., Pfeifer, S., Venn, O., Ségurel, L., Street, T., Leffler, E. M., Bowden, R., Aneas, I., Broxholme, J., Humburg, P., Iqbal, Z., Lunter, G., Maller, J., Hernandez, R. D., Melton, C., Venkat, A., Nobrega, M. A., Bontrop, R., Myers, S., Donnelly, P., Przeworski, M., and McVean, G. (2012). A Fine-Scale Chimpanzee Genetic Map from Population Sequencing. Science, 336:7.

Auton, A. and McVean, G. (2007). Recombination rate estimation in the presence of hotspots. Genome Res., 17(8):1219–1227.

Auton, A., Rui Li, Y., Kidd, J., Oliveira, K., Nadel, J., Holloway, J. K., Hayward, J. J., Cohen, P. E., Greally, J. M., Wang, J., Bustamante, C. D., and Boyko, A. R. (2013). Genetic Recombination Is Targeted towards Gene Promoter Regions in Dogs. PLoS Genet, 9(12):e1003984.

Axelsson, E., Webster, M. T., Ratnakumar, A., Consortium, T. L., Ponting, C. P., and Lindblad-Toh, K. (2012). Death of PRDM9 coincides with stabilization of the recombination landscape in the dog genome. Genome Res., 22(1):51–63.

Baker, C. L., Kajita, S., Walker, M., Saxl, R. L., Raghupathy, N., Choi, K., Petkov, P. M., and Paigen, K. (2015). PRDM9 Drives Evolutionary Erosion of Hotspots in Mus musculus through Haplotype-Specific Initiation of Meiotic Recombination. PLOS Genetics, 11(1):e1004916.

Baker, Z., Przeworski, M., and Sella, G. (2022). Down the Penrose stairs: How selection for fewer recombination hotspots maintains their existence.

Baker, Z., Schumer, M., Haba, Y., Bashkirova, L., Holland, C., Rosenthal, G. G., and Przeworski, M. (2017). Repeated losses of PRDM9-directed recombination despite the conservation of PRDM9 across vertebrates. eLife, 6:e24133.

Baudat, F., Buard, J., Grey, C., Fledel-Alon, A., Ober, C., Przeworski, M., Coop, G., and de Massy, B. (2010). PRDM9 Is a Major Determinant of Meiotic Recombination Hotspots in Humans and Mice. Science, 327(5967):836–840.

Bergeron, L. A., Besenbacher, S., Zheng, J., Li, P., Bertelsen, M. F., Quintard, B., Hoffman, J. I., Li, Z., St. Leger, J., Shao, C., Stiller, J., Gilbert, M. T. P., Schierup, M. H., and Zhang, G. (2023). Evolution of the germline mutation rate across vertebrates. Nature, pages 1–7.

Berglund, J., Quilez, J., Arndt, P. F., and Webster, M. T. (2015). Germline Methylation Patterns Determine the Distribution of Recombination Events in the Dog Genome. Genome Biology and Evolution, 7(2):522–530.

Birney, E., Clamp, M., and Durbin, R. (2004). GeneWise and Genomewise. Genome Res., 14(5):988–995.

Bolívar, P., Mugal, C. F., Nater, A., and Ellegren, H. (2016). Recombination Rate Variation Modulates Gene Sequence Evolution Mainly via GC-Biased Gene Conversion, Not Hill–Robertson Interference, in an Avian System. Mol Biol Evol, 33(1):216–227.

Brick, K., Smagulova, F., Khil, P., Camerini-Otero, R. D., and Petukhova, G. V. (2012). Genetic recombination is directed away from functional genomic elements in mice. Nature, 485(7400):642–645.

Buard, J., Rivals, E., Segonzac, D. D. d., Garres, C., Caminade, P., Massy, B. d., and Boursot, P. (2014). Diversity of Prdm9 Zinc Finger Array in Wild Mice Unravels New Facets of the Evolutionary Turnover of this Coding Minisatellite. PLOS ONE, 9(1):e85021.

Camacho, C., Coulouris, G., Avagyan, V., Ma, N., Papadopoulos, J., Bealer, K., and Madden, T. L. (2009). BLAST+: architecture and applications. BMC Bioinformatics, 10(1):421.

Cavassim, M. I. A., Baker, Z., Hoge, C., Schierup, M. H., Schumer, M., and Przeworski, M. (2022). PRDM9 losses in vertebrates are coupled to those of paralogs ZCWPW1 and ZCWPW2. Proceedings of the National Academy of Sciences, 119(9):e2114401119.

Chan, A. H., Jenkins, P. A., and Song, Y. S. (2012). Genome-Wide Fine-Scale Recombination Rate Variation in Drosophila melanogaster. PLOS Genetics, 8(12):e1003090.

Charlesworth, D., Zhang, Y., Bergero, R., Graham, C., Gardner, J., and Yong, L. (2020). Using GC Content to Compare Recombination Patterns on the Sex Chromosomes and Autosomes of the Guppy, Poecilia reticulata, and Its Close Outgroup Species. Molecular Biology and Evolution, 37(12):3550–3562.

Choi, K., Zhao, X., Tock, A. J., Lambing, C., Underwood, C. J., Hardcastle, T. J., Serra, H., Kim, J., Cho, H. S., Kim, J., Ziolkowski, P. A., Yelina, N. E., Hwang, I., Martienssen, R. A., and Henderson, I. R. (2018). Nucleosomes and DNA methylation shape meiotic DSB frequency in Arabidopsis thaliana transposons and gene regulatory regions. Genome Res., 28(4):532–546.

Clément, Y. and Arndt, P. F. (2013). Meiotic Recombination Strongly Influences GC-Content Evolution in Short Regions in the Mouse Genome. Molecular Biology and Evolution, 30(12):2612–2618.

Davies, B., Hatton, E., Altemose, N., Hussin, J. G., Pratto, F., Zhang, G., Hinch, A. G., Moralli, D., Biggs, D., Diaz, R., Preece, C., Li, R., Bitoun, E., Brick, K., Green, C. M., Camerini-Otero, R. D., Myers, S. R., and Donnelly, P. (2016). Re-engineering the zinc fingers of PRDM9 reverses hybrid sterility in mice. Nature, 530(7589):171–176.

Diagouraga, B., Clément, J. A. J., Duret, L., Kadlec, J., de Massy, B., and Baudat, F. (2018). PRDM9 Methyltransferase Activity Is Essential for Meiotic DNA Double-Strand Break Formation at Its Binding Sites. Molecular Cell, 69(5):853–865.e6.

Duret, L. and Arndt, P. F. (2008). The Impact of Recombination on Nucleotide Substitutions in the Human Genome. PLOS Genetics, 4(5):e1000071.

Eyre-Walker, A. (1999). Evidence of Selection on Silent Site Base Composition in Mammals: Potential Implications for the Evolution of Isochores and Junk DNA. Genetics, 152(2):675–683.

Figuet, E., Ballenghien, M., Romiguier, J., and Galtier, N. (2015). Biased Gene Conversion and GC-Content Evolution in the Coding Sequences of Reptiles and Vertebrates. Genome Biology and Evolution, 7(1):240–250.

Galtier, N. (2021). Fine-scale quantification of GC-biased gene conversion intensity in mammals. Peer Community Journal, 1.

Galtier, N., Roux, C., Rousselle, M., Romiguier, J., Figuet, E., Glémin, S., Bierne, N., and Duret, L. (2018). Codon Usage Bias in Animals: Disentangling the Effects of Natural Selection, Effective Population Size, and GC-Biased Gene Conversion. Molecular Biology and Evolution, 35(5):1092–1103.

Genereux, D. P., Serres, A., Armstrong, J., Johnson, J., Marinescu, V. D., Murén, E., Juan, D., Bejerano, G., Casewell, N. R., Chemnick, L. G., Damas, J., Di Palma, F., Diekhans, M., Fiddes, I. T., Garber, M., Gladyshev, V. N., Goodman, L., Haerty, W., Houck, M. L., Hubley, R., Kivioja, T., Koepfli, K.-P., Kuderna, L. F. K., Lander, E. S., Meadows, J. R. S., Murphy, W. J., Nash, W., Noh, H. J., Nweeia, M., Pfenning, A. R., Pollard, K. S., Ray, D. A., Shapiro, B., Smit, A. F. A., Springer, M. S., Steiner, C. C., Swofford, R., Taipale, J., Teeling, E. C., Turner-Maier, J., Alfoldi, J., Birren, B., Ryder, O. A., Lewin, H. A., Paten, B., Marques-Bonet, T., Lindblad-Toh, K., Karlsson, E. K., and Zoonomia Consortium (2020). A comparative genomics multitool for scientific discovery and conservation. Nature, 587(7833):240–245.

Genestier, A., Duret, L., and Lartillot, N. (2023). Bridging the gap between the evolutionary dynamics and the molecular mechanisms of meiosis: a model based exploration of the PRDM9 intra-genomic Red Queen.

Gerton, J. L. and Hawley, R. S. (2005). Homologous chromosome interactions in meiosis: diversity amidst conservation. Nat Rev Genet, 6(6):477–487.

Glémin, S. (2010). Surprising Fitness Consequences of GC-Biased Gene Conversion: I. Mutation Load and Inbreeding Depression. Genetics, 185(3):939–959.

Glémin, S., Arndt, P. F., Messer, P. W., Petrov, D., Galtier, N., and Duret, L. (2015). Quantification of GC-biased gene conversion in the human genome. Genome Res., 25(8):1215–1228.

Halldorsson, B. V., Hardarson, M. T., Kehr, B., Styrkarsdottir, U., Gylfason, A., Thorleifsson, G., Zink, F., Jonasdottir, A., Jonasdottir, A., Sulem, P., Masson, G., Thorsteinsdottir, U., Helgason, A., Kong, A., Gudbjartsson, D. F., and Stefansson, K. (2016). The rate of meiotic gene conversion varies by sex and age. Nat Genet, 48(11):1377–1384.

Hammoud, S. S., Low, D. H. P., Yi, C., Carrell, D. T., Guccione, E., and Cairns, B. R. (2014). Chromatin and Transcription Transitions of Mammalian Adult Germline Stem Cells and Spermatogenesis. Cell Stem Cell, 15(2):239–253.

Harris, K. and Pritchard, J. K. (2017). Rapid evolution of the human mutation spectrum. eLife, 6:e24284.

He, Y., Wang, M., Dukowic-Schulze, S., Zhou, A., Tiang, C.-L., Shilo, S., Sidhu, G. K., Eichten, S., Bradbury, P., Springer, N. M., Buckler, E. S., Levy, A. A., Sun, Q., Pillardy, J., Kianian, P. M. A., Kianian, S. F., Chen, C., and Pawlowski, W. P. (2017). Genomic features shaping the landscape of meiotic double-strand-break hotspots in maize. Proceedings of the National Academy of Sciences, 114(46):12231–12236.

Hickey, G., Paten, B., Earl, D., Zerbino, D., and Haussler, D. (2013). HAL: a hierarchical format for storing and analyzing multiple genome alignments. Bioinformatics, 29(10):1341–1342.

Hinch, A. G., Zhang, G., Becker, P. W., Moralli, D., Hinch, R., Davies, B., Bowden, R., and Donnelly, P. (2019). Factors influencing meiotic recombination revealed by whole-genome sequencing of single sperm. Science, 363(6433):eaau8861.

Hoge, C. R., Manuel, M. d., Mahgoub, M., Okami, N., Fuller, Z. L., Banerjee, S., Baker, Z., Mcnulty, M., Andolfatto, P., Macfarlan, T. S., Schumer, M., Tzika, A. C., and Przeworski, M. (2023). Patterns of recombination in snakes reveal a tug of war between PRDM9 and promoter-like features.

Kawakami, T., Mugal, C. F., Suh, A., Nater, A., Burri, R., Smeds, L., and Ellegren, H. (2017). Wholegenome patterns of linkage disequilibrium across flycatcher populations clarify the causes and consequences of fine-scale recombination rate variation in birds. Mol Ecol, 26(16):4158–4172.

Kono, H., Tamura, M., Osada, N., Suzuki, H., Abe, K., Moriwaki, K., Ohta, K., and Shiroishi, T. (2014). Prdm9 Polymorphism Unveils Mouse Evolutionary Tracks. DNA Research, 21(3):315–326.

Kumar, S., Suleski, M., Craig, J. M., Kasprowicz, A. E., Sanderford, M., Li, M., Stecher, G., and Hedges, S. B. (2022). TimeTree 5: An Expanded Resource for Species Divergence Times. Molecular Biology and Evolution, 39(8):msac174.

Lartillot, N. (2013). Phylogenetic Patterns of GC-Biased Gene Conversion in Placental Mammals and the Evolutionary Dynamics of Recombination Landscapes. Molecular Biology and Evolution, 30(3):489–502.

Latrille, T., Duret, L., and Lartillot, N. (2017). The Red Queen model of recombination hot-spot evolution: a theoretical investigation. Philosophical Transactions of the Royal Society B: Biological Sciences, 372(1736):20160463.

Lesecque, Y., Glémin, S., Lartillot, N., Mouchiroud, D., and Duret, L. (2014). The Red Queen Model of Recombination Hotspots Evolution in the Light of Archaic and Modern Human Genomes. PLOS Genetics, 10(11):e1004790.

Li, R., Bitoun, E., Altemose, N., Davies, R. W., Davies, B., and Myers, S. R. (2019). A high-resolution map of non-crossover events reveals impacts of genetic diversity on mammalian meiotic recombination. Nat Commun, 10(1):3900.

Lichten, M. and Goldman, A. S. H. (1995). Meiotic recombination hotspots. Annu. Rev. Genet., 29(1):423–444.

Maloisel, L. and Rossignol, J.-L. (1998). Suppression of crossing-over by DNA methylation in Ascobolus. Genes & Development, 12(9):1381–1389.

Marand, A. P., Jansky, S. H., Zhao, H., Leisner, C. P., Zhu, X., Zeng, Z., Crisovan, E., Newton, L., Hamernik, A. J., Veilleux, R. E., Buell, C. R., and Jiang, J. (2017). Meiotic crossovers are associated with open chromatin and enriched with Stowaway transposons in potato. Genome Biol, 18(1):1–16.

Mihola, O., Landa, V., Pratto, F., Brick, K., Kobets, T., Kusari, F., Gasic, S., Smagulova, F., Grey, C., Flachs, P., Gergelits, V., Tresnak, K., Silhavy, J., Mlejnek, P., Camerini-Otero, R. D., Pravenec, M., Petukhova, G. V., and Trachtulec, Z. (2021). Rat PRDM9 shapes recombination landscapes, duration of meiosis, gametogenesis, and age of fertility. BMC Biol, 19(1):1–20.

Mihola, O., Pratto, F., Brick, K., Linhartova, E., Kobets, T., Flachs, P., Baker, C. L., Sedlacek, R., Paigen, K., Petkov, P. M., Camerini-Otero, R. D., and Trachtulec, Z. (2019). Histone methyltransferase PRDM9 is not essential for meiosis in male mice. Genome Res., 29(7):1078–1086.

Milholland, B., Dong, X., Zhang, L., Hao, X., Suh, Y., and Vijg, J. (2017). Differences between germline and somatic mutation rates in humans and mice. Nat Commun, 8(1):15183.

Munch, K., Mailund, T., Dutheil, J. Y., and Schierup, M. H. (2014). A fine-scale recombination map of the human–chimpanzee ancestor reveals faster change in humans than in chimpanzees and a strong impact of GC-biased gene conversion. Genome Res., 24(3):467–474.

Myers, S., Bowden, R., Tumian, A., Bontrop, R. E., Freeman, C., MacFie, T. S., McVean, G., and Donnelly, P. (2010). Drive Against Hotspot Motifs in Primates Implicates the PRDM9 Gene in Meiotic Recombination. Science, 327(5967):876–879.

Nagylaki, T. (1983). Evolution of a finite population under gene conversion. Proceedings of the National Academy of Sciences, 80(20):6278–6281.

Oliver, P. L., Goodstadt, L., Bayes, J. J., Birtle, Z., Roach, K. C., Phadnis, N., Beatson, S. A., Lunter, G., Malik, H. S., and Ponting, C. P. (2009). Accelerated Evolution of the Prdm9 Speciation Gene across Diverse Metazoan Taxa. PLOS Genetics, 5(12):e1000753.

Page, S. L. and Hawley, R. S. (2003). Chromosome Choreography: The Meiotic Ballet. Science, 301(5634):785–789.

Parvanov, E. D., Petkov, P. M., and Paigen, K. (2010). Prdm9 Controls Activation of Mammalian Recom-bination Hotspots. Science, 327(5967):835–835.

Pessia, E., Popa, A., Mousset, S., Rezvoy, C., Duret, L., and Marais, G. A. (2012). Evidence for widespread GC-biased gene conversion in eukaryotes. Genome biology and evolution, 4(7):675–682.

Petes, T. D. (2001). Meiotic recombination hot spots and cold spots. Nat Rev Genet, 2(5):360–369.

Pratto, F., Brick, K., Khil, P., Smagulova, F., Petukhova, G. V., and Camerini-Otero, R. D. (2014). Recombination initiation maps of individual human genomes. Science, 346(6211):1256442–1256442.

Qu, J., Hodges, E., Molaro, A., Gagneux, P., Dean, M. D., Hannon, G. J., and Smith, A. D. (2018). Evolutionary expansion of DNA hypomethylation in the mammalian germline genome. Genome Res., 28(2):145–158.

Quinlan, A. R. and Hall, I. M. (2010). BEDTools: a flexible suite of utilities for comparing genomic features. Bioinformatics, 26(6):841–842.

Raynaud, M., Gagnaire, P.-A., and Galtier, N. (2023). Performance and limitations of linkage-disequilibriumbased methods for inferring the genomic landscape of recombination and detecting hotspots: a simulation study. Peer Community Journal, 3.

Samuk, K. and Noor, M. A. F. (2022). Gene flow biases population genetic inference of recombination rate. G3 Genes|Genomes|Genetics, 12(11):jkac236.

Schield, D. R., Pasquesi, G. I. M., Perry, B. W., Adams, R. H., Nikolakis, Z. L., Westfall, A. K., Orton, R. W., Meik, J. M., Mackessy, S. P., and Castoe, T. A. (2020). Snake Recombination Landscapes Are Concentrated in Functional Regions despite PRDM9. Molecular Biology and Evolution, 37(5):1272–1294.

Singhal, S., Leffler, E. M., Sannareddy, K., Turner, I., Venn, O., Hooper, D. M., Strand, A. I., Li, Q., Raney, B., Balakrishnan, C. N., Griffith, S. C., McVean, G., and Przeworski, M. (2015). Stable recombination hotspots in birds. Science, page 6.

Smagulova, F., Brick, K., Pu, Y., Camerini-Otero, R. D., and Petukhova, G. V. (2016). The evolutionary turnover of recombination hot spots contributes to speciation in mice. Genes Dev., 30(3):266–280.

Smith, T. C. A., Arndt, P. F., and Eyre-Walker, A. (2018). Large scale variation in the rate of germ-line de novo mutation, base composition, divergence and diversity in humans. PLoS Genet, 14(3):e1007254.

Spence, J. P. and Song, Y. S. (2019). Inference and analysis of population-specific fine-scale recombination maps across 26 diverse human populations. Science Advances, 5(10):eaaw9206.

Stapley, J., Feulner, P. G. D., Johnston, S. E., Santure, A. W., and Smadja, C. M. (2017). Variation in recombination frequency and distribution across eukaryotes: patterns and processes. Phil. Trans. R. Soc. B, 372(1736):20160455.

Tock, A. J. and Henderson, I. R. (2018). Hotspots for Initiation of Meiotic Recombination. Front. Genet., 9:521.

Wallberg, A., Glémin, S., and Webster, M. T. (2015). Extreme Recombination Frequencies Shape Genome Variation and Evolution in the Honeybee, Apis mellifera. PLoS Genet, 11(4):e1005189.

Wang, R. J., Thomas, G. W. C., Raveendran, M., Harris, R. A., Doddapaneni, H., Muzny, D. M., Capitanio, J. P., Radivojac, P., Rogers, J., and Hahn, M. W. (2020). Paternal age in rhesus macaques is positively associated with germline mutation accumulation but not with measures of offspring sociability. Genome Res., 30(6):826–834.

Williams, A. L., Genovese, G., Dyer, T., Altemose, N., Truax, K., Jun, G., Patterson, N., Myers, S. R., Curran, J. E., Duggirala, R., Blangero, J., Reich, D., and Przeworski, M. (2015). Non-crossover gene conversions show strong GC bias and unexpected clustering in humans. eLife, 4:e04637.

Wong, W. S. W., Solomon, B. D., Bodian, D. L., Kothiyal, P., Eley, G., Huddleston, K. C., Baker, R., Thach, D. C., Iyer, R. K., Vockley, J. G., and Niederhuber, J. E. (2016). New observations on maternal age effect on germline de novo mutations. Nat Commun, 7(1):10486.

Zelkowski, M., Olson, M. A., Wang, M., and Pawlowski, W. (2019). Diversity and Determinants of Meiotic Recombination Landscapes. Trends in Genetics, 35(5):359–370.

Úbeda, F. and Wilkins, J. F. (2011). The Red Queen theory of recombination hotspots. Journal of Evolutionary Biology, 24(3):541–553.

