## Supplementary_materials for "High prevalence of Prdm9-independent recombination hotspots in placental mammals"

### Supplementary materials from: High prevalence of Prdm9-independent recombination hotspots in placental mammals

November 17, 2023

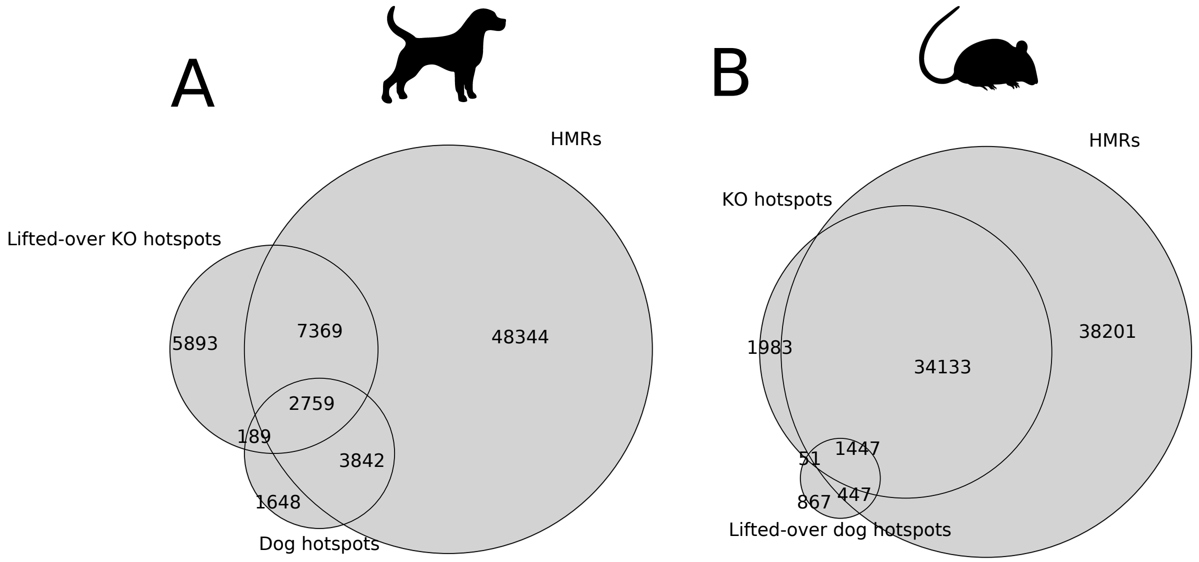


Figure S1: A) Venn diagramm representing the overlap between MDH loci in the dog genome, dog hotspots and dog HMRs in the dog genome. B) Venn diagramm representing the overlap between lifted-over dog hotspots (DRH) on the mouse genome, mouse Prdm9$^{-/-}$ hotspots and mouse HMRs in the mouse genome. Features were considered to overlap if their midpoint were less than 5 kb apart. The numbers indicated in the circles correspond to the number of HMRs that overlap the other features, and dog hotspots in intersections between Prdm9$^{-/-}$ mouse hotspots and dog hotspot.

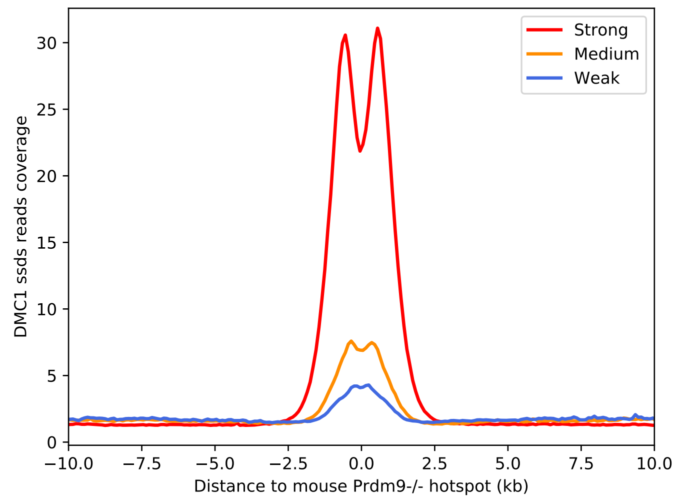


Figure S2: DMC1 ssds read coverage as a function of the distance to the closest MDH in the mouse genome. Hotspots were divided in three equally sized categories of strength: strong hotspots in red (10-190 FPKM), medium hotspots in orange (5-10 FPKM) and weak hotspots in blue (0-5 FPKM). The line directly correspond to the mean value of DMC1 ssds read coverage in a 100 bp window.

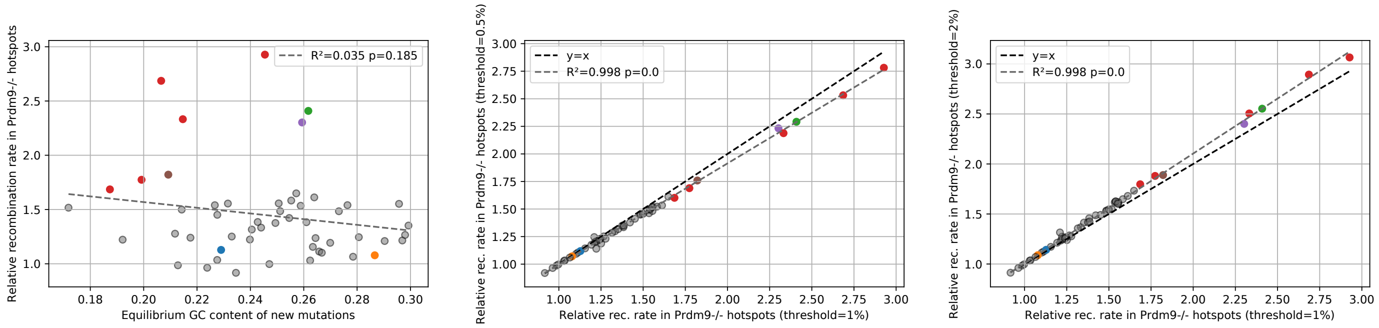


Figure S3: A) Relative recombination rate in lifted-over mouse Prdm9$^{-/-}$ hotspots as a function of the estimated $GC^*$ of new mutations in all 52 species of mammals. B) and C) Relative recombination rate in lifted-over mouse Prdm9$^{-/-}$ hotspots using different thresholds to estimate $GC^*_{mut}$ (0.5% in B and 2% in C) as a function of the relative recombination rate in lifted-over mouse Prdm9$^{-/-}$ hotspots using the 1% threshold used throughout the study in all 52 species of mammals. Red points correspond to canids, the green point indicate ring-tailed lemurs, the purple point northern elephant seals, the blue point mice, the brown point daurian ground squirrels, and the orange point humans.

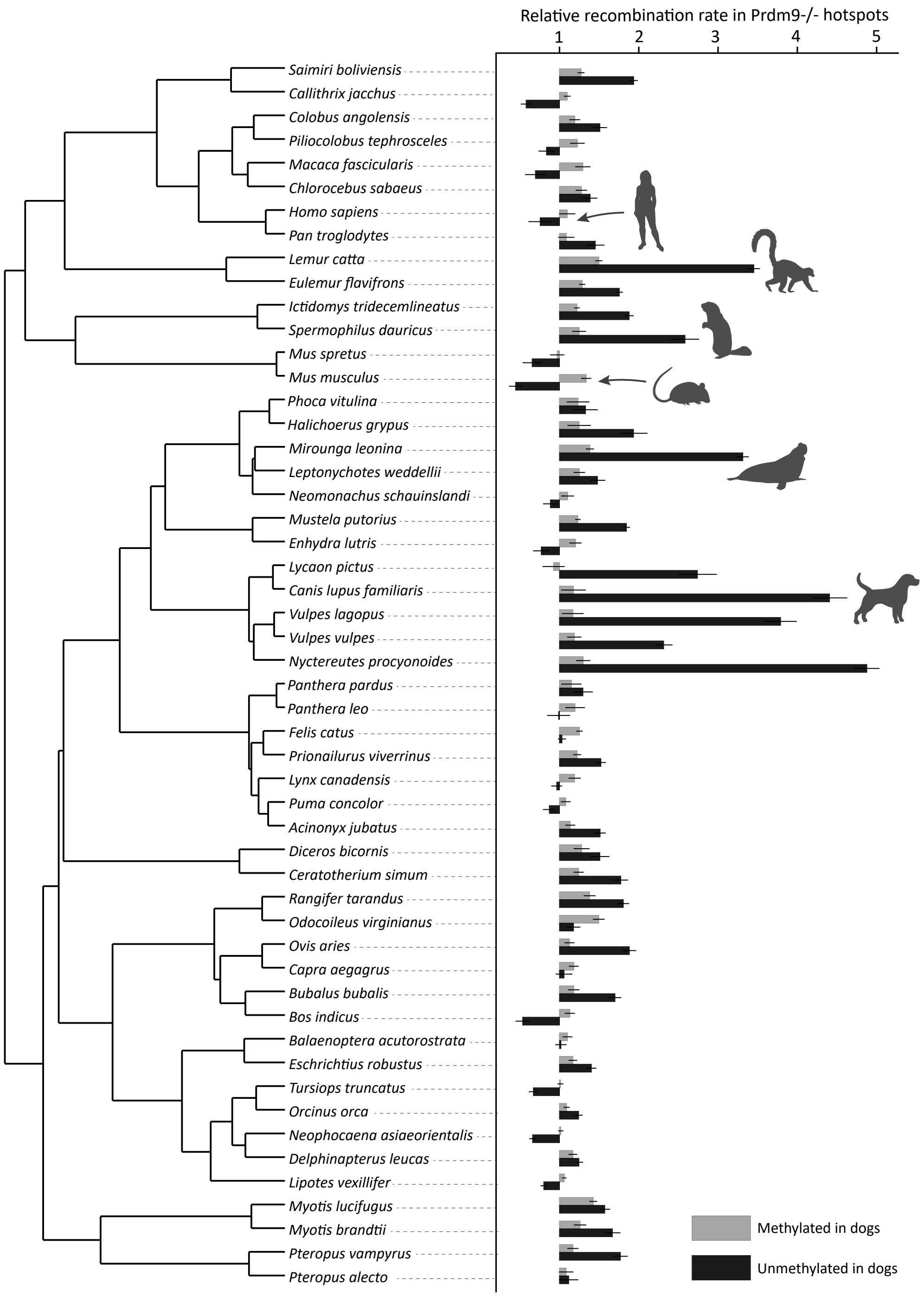


Figure S4: Relative recombination rates at MDH loci in 52 mammals. The hotspots were splited in two categories depending on their methylation level in dog sperm. The number of lifted features for each categories varied from ∼4,000 in *Myotis brandtii* to ∼9,000 in *Mus spretus*. Detailed numbers of mapped features are presented in Supplementary Table XX. The tree have been retrieved from TimeTree5 Kumar et al. (2022). Error bars correspond to a 95% confidence interval obtained by bootstrapping the substitutions for computing $GC^*_{flank}$ and $GC^*_{hot}$.

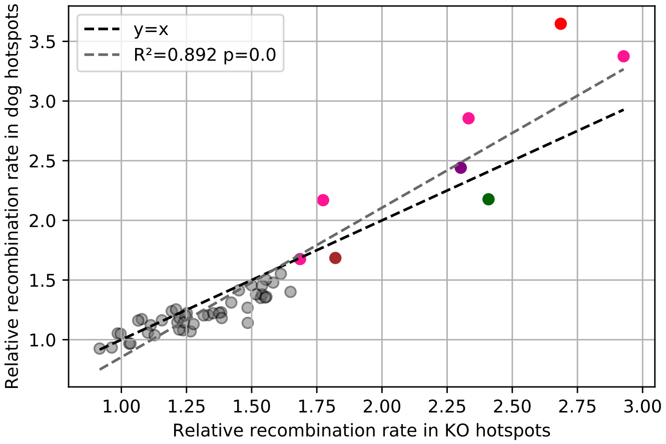


Figure S5: Relative recombination rate in lifted-over dog LD-based hotspots as a function of the relative recombination rate in MDH loci. Red points correspond to canids, the green point indicate ring-tailed lemurs, the purple point northern elephant seals, the blue point mice, the brown point daurian ground squirrels, and the orange point humans.

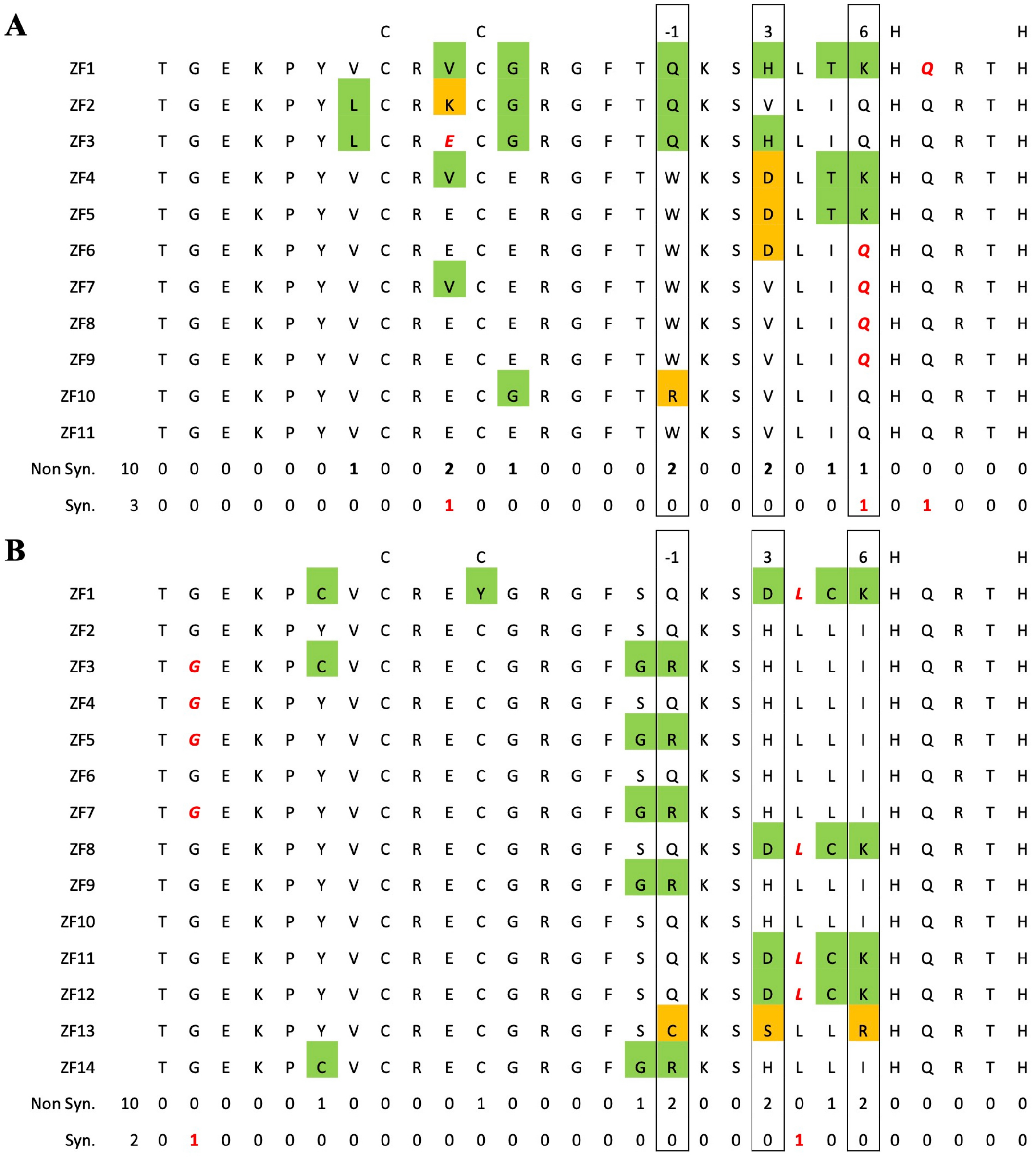


Figure S6: Sequence diversity of zinc finger arrays of the Prdm9 allele present in the reference genome of *Spermophilus dauricus* (A) and *Lemur catta* (B). Amino-acid variants are highlighted by a colored background. Synonymous variants are written in red. The number of non-synonymous and synonymous variants is indicated at each position. In both species, amino-acid variants are enriched at positions -1, 3, and 6 that are involved in DNA binding. Under neutral evolution, one would expect approximatively one third of variants to be synonymous and two thirds to be non-synonymous. In both species, the number of non-synonymous variants exceeds the neutral expectation, which is suggestive of positive selection. *Lemur catta* Prdm9: RefSeq protein accession number: XP_045389275.1_1, genomic coordinates of the exon encoding the zinc finger array: complement(NC_059147.1: 34824301..34826009). *Spermophilus dauricus* Prdm9: the protein is not annotated in RefSeq, genomic coordinates of the exon encoding the zinc finger array: KZ296155.1: 118252..119699.

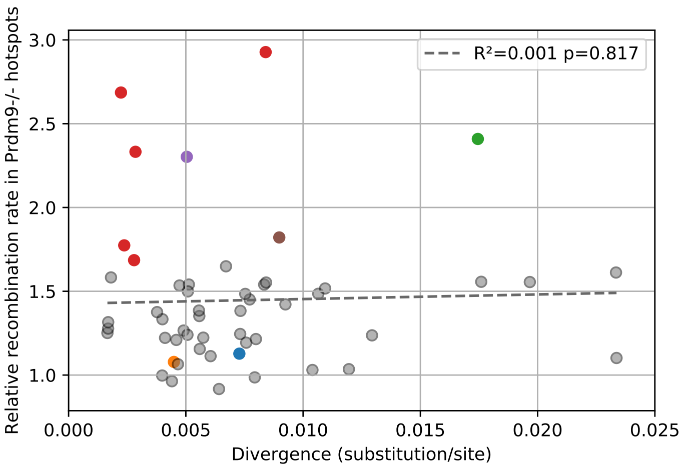


Figure S7: Relative recombination rate in lifted-over mouse Prdm9$^{-/-}$ hotspots as a function of the estimated length of the branch used to compute $GC^*$ in substitutions per/site. Total number of sites were computed with sites for which all threes species (target sister and outgroup) were genotyped and aligned, excluding CpG dinucleotides. Red points correspond to canids, the green point indicate ring-tailed lemurs, the purple point northern elephant seals, the blue point mice, the brown point daurian ground squirrels, and the orange point humans.

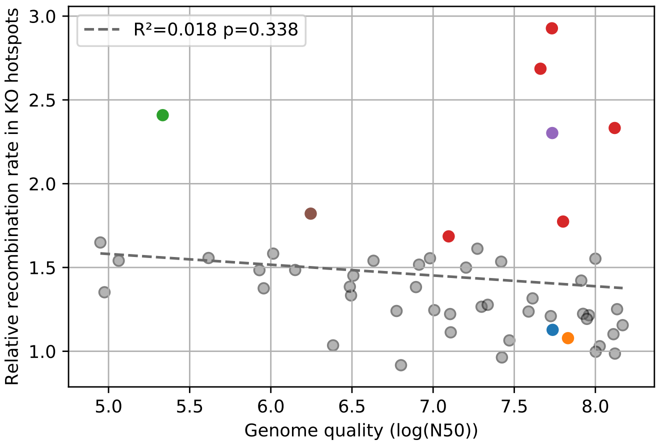


Figure S8: Relative recombination rate in lifted-over mouse Prdm9$^{-/-}$ hotspots as a function of the genome quality summarized by the log of the N50 metric. Red points correspond to canids, the green point indicate ring-tailed lemurs, the purple point northern elephant seals, the blue point mice, the brown point daurian ground squirrels, and the orange point humans.

### References

Sudhir Kumar, Michael Suleski, Jack M Craig, Adrienne E Kasprowicz, Maxwell Sanderford, Michael Li, Glen Stecher, and S Blair Hedges. TimeTree 5: An Expanded Resource for Species Divergence Times. *Molecular Biology and Evolution*, 39(8):msac174, August 2022. ISSN 1537-1719. doi: 10.1093/molbev/msac174. URL https://doi.org/10.1093/molbev/msac174.
